# Elevational Shifts in Tropical Tree Leaf Traits: Interactions Between Soil, Climate, Light, and Phylogeny

**DOI:** 10.1101/2025.07.30.667609

**Authors:** Jiří Doležal, Kirill Korznikov, Vaclav Bazant, Adam Taylor Ruka, Jorge Gago

## Abstract

Understanding how tropical trees respond to complex environmental gradients is essential for predicting forest resilience under climate change. We examined variation in key leaf traits, including specific leaf area (SLA), foliar nitrogen (N) and phosphorus (P), C:N and N:P ratios, and stable isotope composition (δ^13^C, δ^15^N), in 160 tree species along a 3,200-m elevational transect on Mount Cameroon. This gradient spans hyper-humid coastal rainforests to arid Afroalpine savannas, capturing sharp transitions in climate, soils, and forest structure. Leaf traits shifted nonlinearly with elevation, from acquisitive strategies at mid-elevations to conservative syndromes in lowlands and highlands. Mid-elevation forests (∼1,000–1,500 m), characterized by moderate climate and canopy disturbance by elephants, supported nutrient-rich, high-SLA foliage. In contrast, high-elevation forests (>2,000 m) exhibited low SLA, high C:N, and enriched δ^13^C, consistent with stress tolerance under cold, dry, and fire-prone conditions. The strongest phosphorus limitation occurred in hyper-humid lowlands, where extreme rainfall (>12,000 mm/year) drives leaching losses. Foliar δ^15^N declined markedly with elevation (from +5‰ to −5‰), indicating a shift from mineral N uptake and N-fixation in lowland Fabaceae to ecto- and ericoid mycorrhizal associations in montane Ericaceae. A bimodal δ^15^N pattern, with enrichment in both lowland and upper montane forests, reflects N-fixation under leaching and fire-driven N scarcity, respectively. Phylogenetic analyses showed that climate, soils, forest structure, and lineage jointly shaped trait–environment relationships. Traits related to δ^13^C, C:N, and δ^15^N exhibited strong phylogenetic signal, highlighting evolutionary constraints. These findings underscore the value of integrating functional traits, isotopes, and phylogeny to predict tropical forest responses to global change.

## 1. Introduction

Understanding how plant functional traits vary across environmental gradients is central to ecology, biogeography, and global change biology (Wright et al., 2004; Reich, 2014; Garnier et al., 2015). Leaf traits such as specific leaf area (SLA), leaf nitrogen (N) and phosphorus (P) content, and leaf N:P ratios are widely used as proxies for resource acquisition strategies, photosynthetic capacity, and growth responses (Wright et al., 2004; Reich, 2014). These traits are shaped by multiple environmental filters, including nutrient availability, moisture regimes, temperature, and light conditions, that often covary along elevation gradients (Körner, 2021). However, because these drivers are interdependent, it can be challenging to isolate their respective contributions to trait variation, which limits the ecological interpretation and predictive utility of trait-based approaches (Fyllas et al., 2017; Oliveras et al., 2020). Despite the rise of global trait databases, tropical mountain systems, particularly those in Africa, remain underrepresented in trait-based ecology, despite sustaining high biodiversity and regulating major global carbon and water fluxes.

Elevational gradients provide natural laboratories for examining trait–environment relationships, encompassing steep and relatively continuous transitions in climate, edaphic conditions, and forest structure over short geographic distances (Körner, 2007; Girardin et al., 2014). While such gradients have been widely used in temperate and boreal regions, studies spanning tropical rainforest to high-montane savanna remain scarce. The elevational transect on Mount Cameroon, a geologically young, exceptionally steep volcano rising from sea level to over 4,000 m, is one of the most dramatic tropical elevation gradients globally (Hořák et al., 2019; Maicher et al., 2020; Doležal et al., 2023). Yet, functional trait data from this region are virtually absent from global syntheses. This setting presents a rare opportunity to investigate how plant strategies transition from humid lowland rainforest to montane and Afroalpine ecosystems, thereby filling a critical empirical gap in trait-based biogeography and ecosystem modeling.

In tropical forest systems, lowland forests are generally characterized by high temperatures, relatively stable moisture conditions, and high radiation, while montane forests experience cooler temperatures, frequent cloud immersion, and often lower light availability (Kitayama & Aiba, 2002; Girardin et al., 2016). Soil fertility may decline or shift in nutrient composition with elevation, depending on parent material, erosion, and leaching processes (Vitousek & Sanford, 1986; Soethe et al., 2008). Leaf trait variation along these gradients, such as declining SLA or leaf N content with elevation, is often interpreted as an adaptive response to colder or nutrient-limited conditions. Yet such patterns may also reflect light limitation under dense canopies or nutrient reallocation in response to drought or other environmental stressors.

Distinct environmental factors constrain different tropical forests, and these constraints are reflected in characteristic leaf trait syndromes (Pringle et al., 2011; Méndez-Alonzo et al., 2012). In seasonally dry tropical forests or savanna ecotones, where water availability is the principal limiting factor, plants tend to exhibit conservative leaf traits, such as low SLA, and low leaf N and P concentrations, associated with drought tolerance and slow resource turnover (Markesteijn & Poorter, 2009; Wright et al., 2001). By contrast, in moist lowland rainforests where light is the main limiting resource beneath a closed canopy, species often display higher SLA and elevated leaf nutrient content, reflecting acquisitive strategies optimized for high light capture and fast growth in gaps or upper canopy strata (Díaz et al., 2016; Asner et al., 2018). In tropical montane or cloud forests, low temperatures and persistent cloud cover can limit both photosynthesis and transpiration, leading to thick, tough leaves (low SLA, high LDMC), relatively low nutrient concentrations, and in some cases, decoupling of leaf traits from typical stoichiometric expectations due to constrained nutrient uptake or translocation (Vitousek et al., 1992; Moser et al., 2007). In nutrient-poor forests, such as those developed on old or highly weathered tropical soils, species often exhibit low leaf nutrient concentrations and high N:P ratios, indicating phosphorus limitation (Townsend et al., 2007; Porder & Hilley, 2011).

Leaf traits can thus serve as ecological indicators of the dominant resource constraints in a given habitat. High SLA and high leaf N are commonly associated with nutrient-rich and high-light environments, whereas low SLA, high LDMC, and low leaf nutrient content are indicative of stress tolerance under drought, cold, or nutrient limitation (Wright et al., 2004; Laughlin, 2014). Leaf N:P ratios provide further insights into the type of nutrient limitation: low N:P (<10–14) suggests nitrogen limitation, high N:P (>16–20) indicates phosphorus limitation, and intermediate values suggest co-limitation or balanced nutrition (Koerselman & Meuleman, 1996; Güsewell, 2004). These trait–environment relationships are increasingly being used to infer ecosystem-level nutrient cycling, plant resource strategies, and responses to environmental change.

Stable isotope ratios of carbon (δ^13^C) and nitrogen (δ^15^N) in leaves provide an additional layer of inference, reflecting the integration of physiological and ecological processes (Givnish, 2002; Santiago et al., 2017). Leaf δ^13^C is widely used as a proxy for intrinsic water-use efficiency (WUEi), as it reflects the ratio of carbon assimilation to stomatal conductance over time (Farquhar et al., 1989). Higher δ^13^C values indicate greater WUE and are often associated with plants experiencing drought stress or growing in arid microclimates (Cernusak et al., 2007; Cornwell et al., 2018). In elevational gradients, δ^13^C typically increases (becomes less negative) with altitude by about 1.1‰ per kilometer on average, due to reduced atmospheric pressure and CO_2_ partial pressure (Körner et al., 1991). However, it can also vary with canopy position and light exposure, as well as increasing evaporative demand (Holtum & Winter, 2005; Doležal et al., 2021). As leaves become thicker and denser with elevation, as reflected in decreasing SLA, a corresponding decline in leaf nutrient content and increased δ^13^C may suggest a trade-off between photosynthetic nitrogen efficiency and water-use efficiency (Wright et al., 2001; Cornwell et al., 2018). Thicker leaves, while more durable and better protected against abiotic stressors, often limit CO_2_ diffusion (mesophyll conductance), resulting in lower photosynthetic capacity per unit N and higher δ^13^C values. Thus, a negative correlation between leaf N and δ^13^C across the gradient could be expected, and may reflect this fundamental trade-off in resource allocation in response to increasingly harsh abiotic conditions and reduced competitive pressure at the treeline.

In contrast, δ^15^N values may exhibit a complex pattern, diverging from δ^13^C trends and necessitating a more nuanced interpretation. δ^15^N is influenced by N cycling processes, symbiotic associations, and rooting depth (Craine et al., 2015; Yang et al., 2015). Mycorrhizal associations, especially with ectomycorrhizal or ericoid fungi, tend to result in lower δ^15^N values due to fractionation during uptake (Högberg, 1997; Mayor et al., 2014). Conversely, Fabaceae often show higher or near-zero values, depending on their N source (e.g., symbiotic fixation, nitrate vs. ammonium uptake). δ^15^N values may thus reflect both ecological processes (e.g., nutrient access, waterlogging, or periodic anoxia at low elevation) and phylogenetic filtering along the gradient, underscoring the need for careful disentangling of abiotic drivers from lineage-specific trait syndromes (Cornwell & Ackerly, 2009; Laughlin & Messier, 2015).

In this study, we investigate elevational trends in key leaf traits across 160 tropical tree species sampled along a nearly 3,200 m elevational gradient on Mount Cameroon, spanning from sea-level rainforests to upper montane and Afroalpine vegetation. For each species, we measured SLA, leaf N and P content, N:P ratios, and stable isotope composition (δ^13^C and δ^15^N), alongside structural traits (tree height, stem diameter) and detailed environmental data, including soil chemistry, microclimate (temperature, humidity), and light availability (from hemispherical photography). We hypothesized that trait variation across the gradient reflects a combination of environmental filtering and evolutionary conservatism, with acquisitive strategies (e.g., high SLA, high leaf N and P, low δ^13^C) favored in warm, fertile lowlands, and conservative traits (e.g., low SLA, high C:N, elevated δ^13^C) emerging in colder, drier, and nutrient-limited high elevations. These predictions are grounded in the leaf economic spectrum (LES) framework, which posits that plant traits are structured by trade-offs between resource acquisition and conservation. However, we also expected deviations from LES predictions in areas where fire, hydrology, forest elephant disturbances, or nutrient decoupling disrupt classical trait–soil patterns. By testing these hypotheses, we aim to identify the dominant environmental constraints underlying observed trait variation and to advance trait-based ecological frameworks for Afrotropical forests, one of the world’s most under-sampled and climatically dynamic ecosystems.

### 2.1. Study Area

This study was conducted along a continuous elevational gradient on the southwestern slopes of Mount Cameroon, located in the Republic of Cameroon (Figure 1). Data were collected from 176 permanent monitoring plots spanning a near-pristine rainforest-to-alpine gradient, ranging from coastal lowland forests to the treeline at approximately 3200 m a.s.l., a unique ecological continuum in West Africa, situated within Mount Cameroon National Park (Maicher et al. 2020). The plots encompassed all major vegetation zones, including forested areas from sea level to montane cloud forests at 2400 m, as well as isolated tree stands and solitary trees extending into the afromontane savanna up to 3200 m (Dolezal et al., 2022). At each elevation level, 16 plots were systematically established approximately 150 meters apart along the contour to capture variability in vegetation structure, canopy light conditions, leaf functional traits, and soil chemistry. Because the southwestern transect of the mountain is bordered by agricultural land and coastal settlements below 300 m a.s.l., the lowland rainforest was instead sampled within the nearby Bimbia Forest Reserve. This protected coastal forest harbors high tree species richness, with over 100 species per hectare, and provides a representative baseline for the coastal rainforest zone (Ferenc et al., 2018; Maicher et al., 2020).

**Figure 1.**
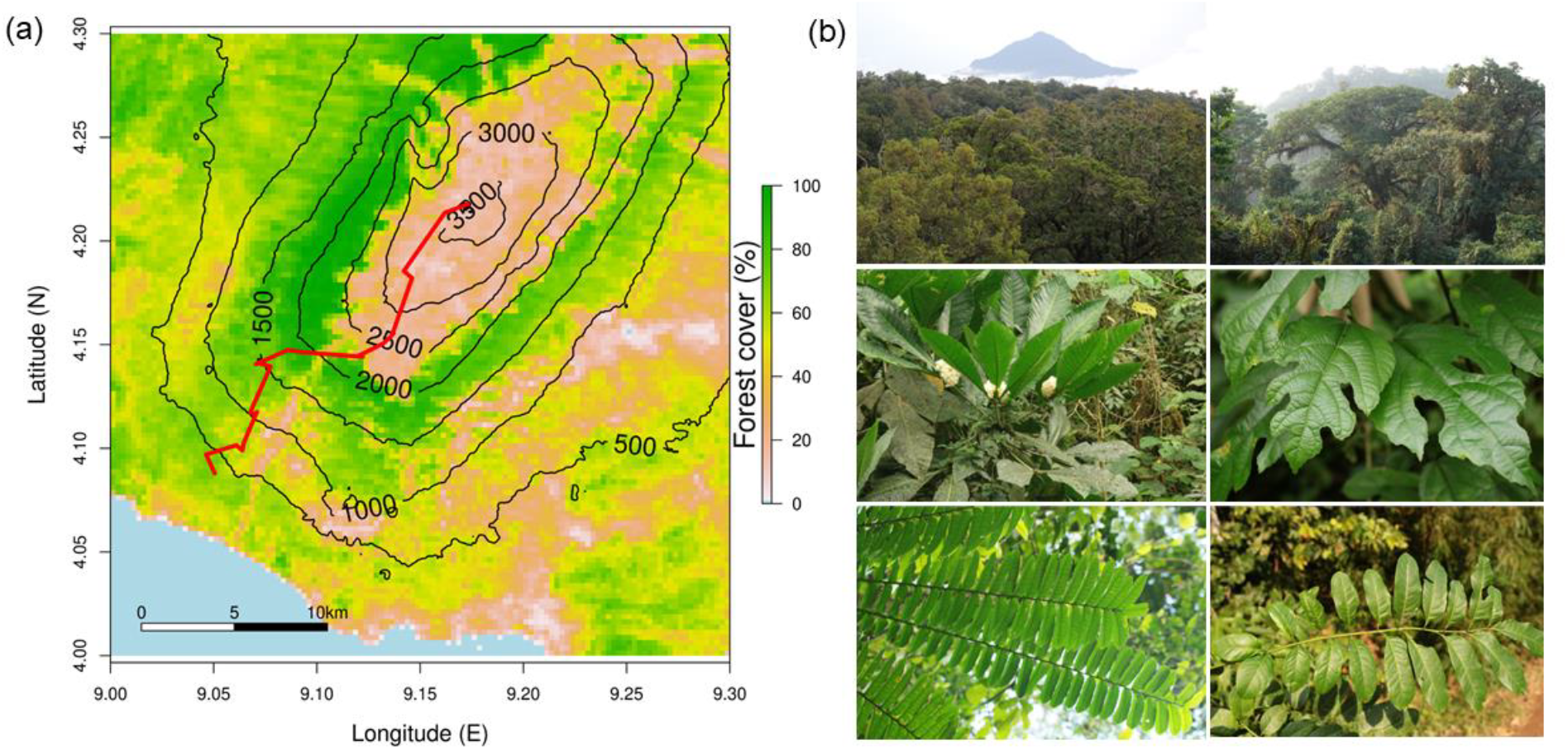
A study area on the southwestern slopes of Mount Cameroon, West Africa. (a) The map shows forest cover (%) with red transects indicating sampling locations. (b) Photos on the right illustrate vegetation types and leaf morphology along the gradient.

Mount Cameroon lies within a perhumid tropical climate zone influenced by seasonal shifts between the moist southwest monsoon and the dry northeast harmattan winds. The region experiences a distinct rainy season from June to September and a dry season from late December to February, with brief transitional periods in between. Annual precipitation in the coastal lowlands exceeds 11,500 mm (Rehakova et al., 2022), making this one of the wettest locations on the Planet. Rainfall decreases steadily with increasing elevation and distance from the ocean, creating a pronounced dry belt between 2300 and 3200 m. This rain shadow effect facilitates the development of a treeless afromontane savanna above the forest limit (Dolezal et al. 2022).

The flora of Mount Cameroon is exceptionally diverse, with more than 2300 vascular plant species recorded, encompassing over 800 genera and 210 families (Cheek et al. 1996; Cable and Cheek 1998). The mountain’s forest vegetation is stratified into distinct zones along the elevational gradient (Hall, 1973; Payton, 1993; Proctor et al., 2007). Coastal rainforest (0– 250 m a.s.l.) is dominated by tall *Desbordesia glaucescens*, along with *Ceiba pentandra, Coelocaryon preussii, Pycnanthus angolensis, Scyphocephalium mannii*, and *Lophira alata*. A closed canopy and sparse understory characterize this zone. Lowland rainforest (300–1000 m a.s.l.) harbors species such as *Pycnanthus angolensis, Coelocaryon preussii, Crudia gabonensis, Berlinia bracteosa, Anthonotha fragrans*, and *Baikiaea insignis*, characterized by a structurally intact canopy. Mid-elevation forest (1000–1600 m a.s.l.) forms a transitional belt marked by reduced canopy closure due to disturbance by forest elephants (*Loxodonta cyclotis*), which maintain a mosaic of open clearings and closed forest. Dominant species include *Anthonotha noldeae, Schefflera mannii, Ficus chlamydocarpa*, and *Entandrophragma utile*. Montane forest (1600–2300 m a.s.l.) occurs under persistent cloud cover and mist, with a denser understory and canopy dominated by *Ilex mitis, Schefflera mannii, S. abyssinica, Syzygium staudtii*, and *Prunus africana*. Afromontane vegetation (2400–2300 m a.s.l.) is dominated by C4 grasses (*Loudetia simplex, Pennisetum monostigma*) and forbs (*Trifolium simense, Satureja robusta*), with sparse stands of fire-tolerant trees such as *Myrica arborea* and *Agarista salicifolia*, limited by volcanic activity, porous substrate, and low precipitation. Although the climatic tree line would occur around 3200–3500 m based on temperature, actual tree growth ceases at much lower elevations. At the highest elevations, from approximately 3300 m to the summit at 4000 m, the Afroalpine zone is characterized by cold, windy conditions and sparse plant cover, composed of C3 grasses, herbs, and low-growing shrubs (Dolezal et al., 2022).

### 2.2. Leaf Trait Sampling

We analyzed leaf functional trait data from 412 individual trees collected across a network of 176 monitoring plots, representing 164 species, 122 genera, and 53 families (Fig. 2), along an elevational gradient on Mount Cameroon. We used a single rope technique and 3-m-long pruning shears to collect 15–20 leaves from the upper, well-lit crown regions. Fresh leaf blades were photographed in the field to assess leaf area and SLA. For each individual, we measured eight leaf traits: leaf area (LA), specific leaf area (SLA), leaf nitrogen (LNC), leaf phosphorus (LPC), leaf carbon (LCC) content, carbon-to-nitrogen ratio (C:N), nitrogen-to-phosphorus ratio (N:P), and stable isotope ratios of carbon (δ^13^C) and nitrogen (δ^15^N). Total carbon and nitrogen content, along with δ^13^C and δ^15^N, were measured at the Stable Isotope Facility at UC Davis, USA, using an elemental analyzer-isotope ratio mass spectrometer (EA-IRMS). Phosphorus concentrations were determined via spectrophotometry following HClO_4_ digestion using a SHIMADZU UV-1650PC spectrophotometer.

**Figure 2.**
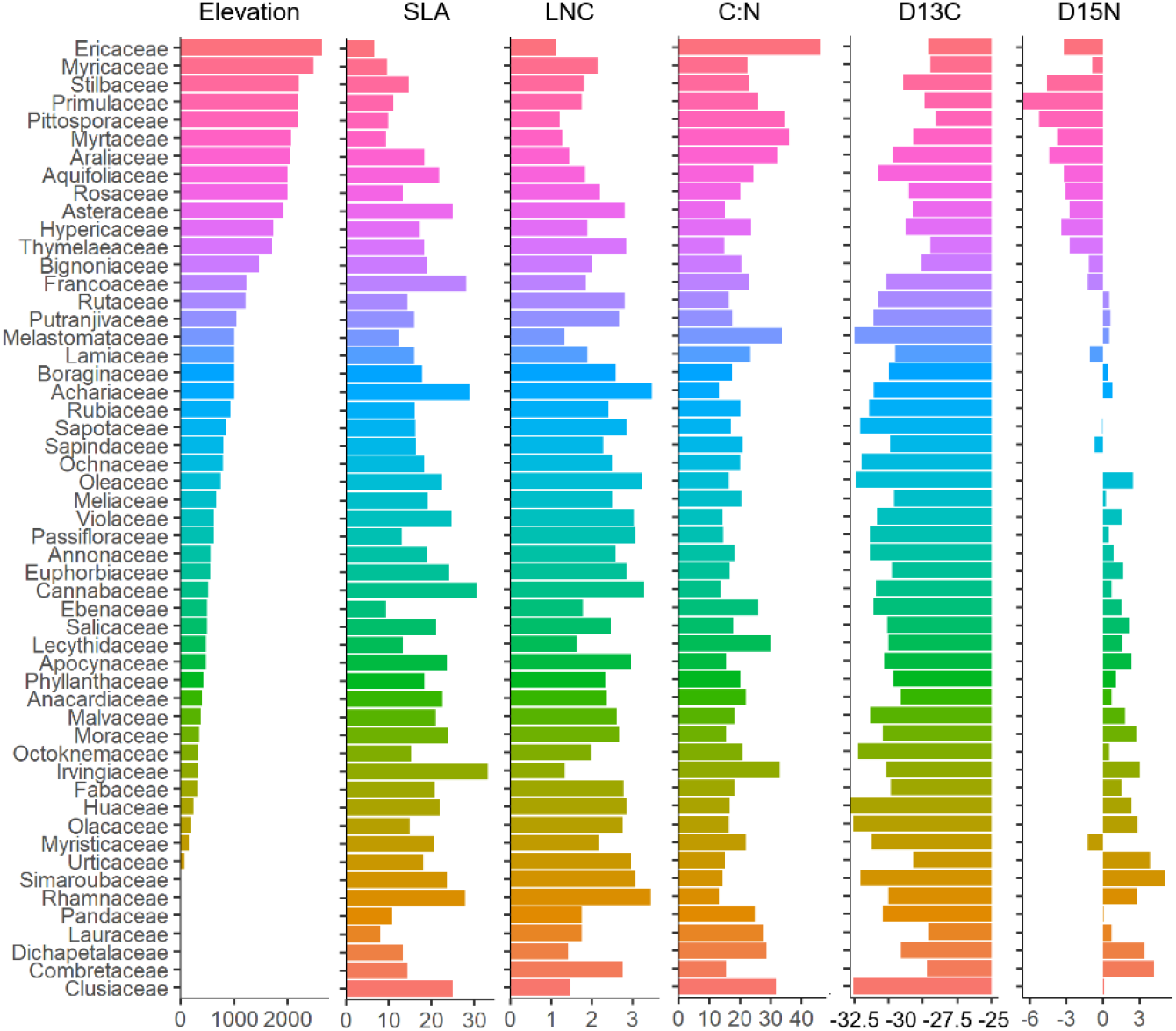
Variation in leaf traits across plant families along an elevational gradient on the SW slopes of Mount Cameroon. Bars represent family means for elevation, SLA, LNC, foliar C:N ratio, stable carbon isotope ratio (δ^13^C), and nitrogen isotope ratio (δ^15^N). Families are ordered by mean elevation, from high-elevation taxa (e.g., Ericaceae, Primulaceae, Myricaceae) to low-elevation taxa (e.g., Combretaceae, Clusiaceae), highlighting coordinated trait shifts from stress tolerance to resource acquisitiveness.

### 2.3. Vegetation Structure and Light Environment

To assess the relationship between leaf traits, forest structure, and species composition, we conducted a detailed inventory of all trees with a diameter at breast height (DBH) exceeding 10 cm in each plot. Each tree was georeferenced, identified to the species level, and measured for DBH and total height. In total, we recorded over 10,000 individual trees belonging to more than 250 species. From these measurements, we derived several structural metrics for each forest plot, including stand basal area, total volume of live and dead trees, mean and maximum stem diameter (DBH) and tree height, the coefficients of variation (CV) for DBH and height (to capture vertical stratification), and the stem slenderness index (SSI). Canopy light conditions were quantified using hemispherical photography. In each forest plot, five hemispherical images were captured at a height of 1.8 meters aboveground, one at the plot center and four at points 10 meters away in the cardinal directions, resulting in a total of 480 photographs. These images were analyzed with WinSCANOPY software to estimate canopy openness, leaf area index, and both direct and diffuse solar radiation penetration.

### 2.4. Climate Variables

Due to the absence of meteorological stations above 1000 m a.s.l. on Mount Cameroon, we relied on high-resolution gridded climate data from the TerraClimate database (Abatzoglou et al., 2018) to characterize the climatic conditions across our study sites. This dataset provides monthly records of multiple climate variables, including mean air temperature, total precipitation, vapor pressure deficit (VPD), and the Palmer Drought Severity Index (PDSI), at a spatial resolution of 1/24 degrees. For each monitoring plot, we extracted the corresponding grid cell data and calculated the long-term mean annual values for the period from 2000 to 2014. To ground-truth the accuracy of these gridded estimates and ensure they reflect the microclimatic realities of the elevational gradient on Mount Cameroon, we conducted extensive field-based climate monitoring. Approximately 160 temperature dataloggers (EMS Brno and TOMST Praha, Czechia) were deployed at hourly resolution along the full elevational transect. Additionally, we installed 15 manual rain gauges (EMS Brno, Czechia) at key sites to capture site-specific precipitation data. These gauges were regularly maintained to remove accumulated vegetation, leaf litter, and other debris, and were protected from damage, particularly from seasonal fires in the savanna zone. Comparative analyses revealed strong correlations between the gridded climate data and the in situ measurements, with Pearson correlation coefficients ranging from 0.7 to 0.9 (P < 0.01). This high degree of agreement validates the use of the TerraClimate dataset as a reliable representation of local climate conditions along Mount Cameroon’s complex elevational gradient.

### 2.5. Soil characteristics

To assess soil fertility, approximately 150 g of soil was collected from each plot. Samples were oven-dried at 100 °C, manually ground using a mortar, and then sieved through a 2 mm mesh after removing coarse roots and organic debris (Rehakova et al., 2022). Key macronutrients, including calcium (Ca^2^_+_), magnesium (Mg^2^_+_), potassium (K_+_), sodium (Na_+_), nitrogen, and phosphorus, were analyzed for each sample (see Supplementary Table S1). Cation concentrations were determined using atomic absorption spectroscopy (SpectrAA 640, Varian Techtron). Nitrogen forms (ammonium, nitrate, and total nitrogen) were measured colorimetrically following Kjeldahl digestion, using a FIAstar 5010 flow injection analyzer (Tecator). Phosphorus was quantified colorimetrically after perchloric acid (HClO_4_) digestion using a SHIMADZU UV-1650PC spectrophotometer. In addition to nutrient concentrations, we assessed a suite of basic soil properties, including pH, gravimetric water content, soil carbon content (SOC), and soil texture, expressed as the proportion of coarse particles (diameter>0.5 mm) (Fig. 3).

**Figure 3.**
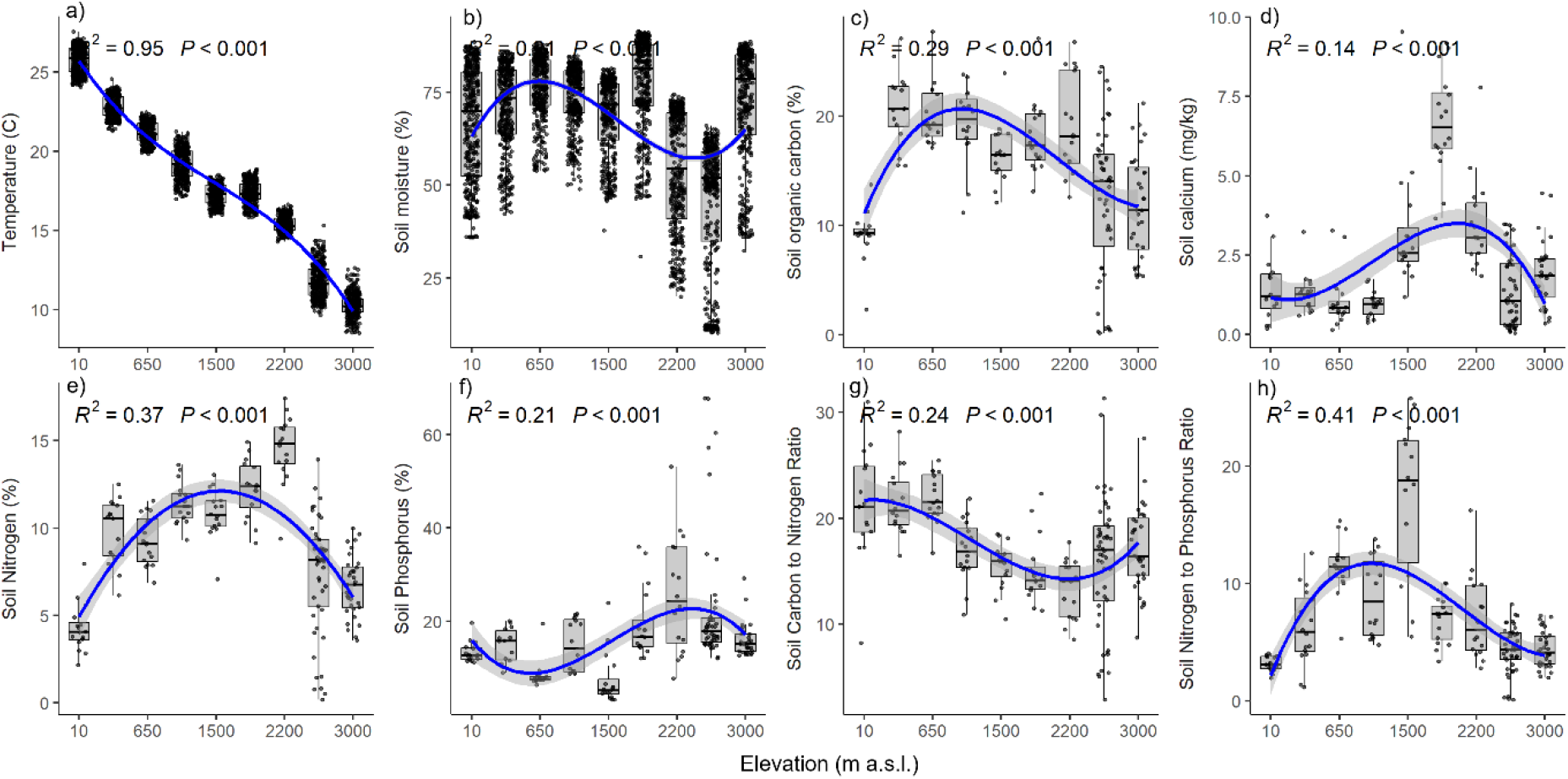
Environmental variation along the elevational gradient. Relationships between (a) elevation and temperature, (b) soil moisture, and soil chemical properties, including (c) organic carbon, (d) calcium, (e) nitrogen, (f) phosphorus, (g) C:N ratio, and (h) N:P ratio. Blue lines indicate fitted GAM (generalized additive model) curves; shaded areas show 95% confidence intervals. Boxplots represent binned elevation intervals. All relationships are statistically significant (P < 0.001).

### 2.6. Data analysis

We used generalized additive models (GAM) to determine changes in environmental parameters (climate, soil, forest structure) and leaf traits along the elevational gradient studied (Fig. 3). The GAM framework helps discover elevation-specific patterns, does not rely on any predefined relationships, and provides easy-to-interpret visualizations (McCullagh and Nelder 1989). Because many leaf traits were intercorrelated, we conducted a principal component analysis (PCA) to summarize the major axes of variation in these traits (Fig. 4). Leaf trait GAMs were fitted for all individuals and separately for small-statured species (typically subcanopy trees <15 m tall) and large-statured species (canopy and emergent trees) (Fig. 5). To assess how environmental factors, encompassing climate, soil, light, and forest structure, shape functional trait variation across tropical montane tree species, we quantified phylogenetically-informed standardized effect sizes (SES) for associations between nine key leaf traits and 30 environmental variables encompassing climate, soil, and forest structure. To evaluate the extent to which trait–environment relationships reflect shared evolutionary history, we quantified phylogenetic signal using standardized effect sizes (SES) derived from a phylogenetic permutation framework. For each trait–environment pair, we compared the observed correlation coefficient to a null distribution generated by 999 random tip-shufflings of the phylogeny, which preserves trait and environmental values but disrupts evolutionary structure. Significant positive SES values (|SES| > 2) indicate that closely related species tend to respond similarly to a given environmental gradient, consistent with phylogenetic clustering and trait conservatism. For example, a strong positive association between foliar δ^15^N and mean annual temperature (MAT) would suggest that lineages adapted to warmer environments, such as Fabaceae or Euphorbiaceae, consistently exhibit higher δ^15^N values. In contrast, significant negative SES values indicate weaker-than-expected trait–environment coupling under phylogenetic constraint, suggesting that phylogenetic overdispersion may arise from convergent evolution or ecological equivalence across distantly related taxa. The phylogenetic distance matrix was constructed from a species-level megaphylogeny using patristic distances. Leaf trait data (e.g., specific leaf area, nitrogen, δ^13^C) were averaged across individuals per species. Environmental variables were extracted from plot-level soil and light measurements or interpolated climate surfaces and averaged by species occurrence. Analyses were conducted in Python using numpy, scipy, pandas, and matplotlib. Phylogenetic permutations and SES computation were implemented with custom scripts. We reconstructed the phylogeny of the studied species using MrBayes 3.1.2, based on nucleotide sequences obtained from GenBank and those generated from field-collected material (Plavcova *et al*., 2024).

**Figure 4.**
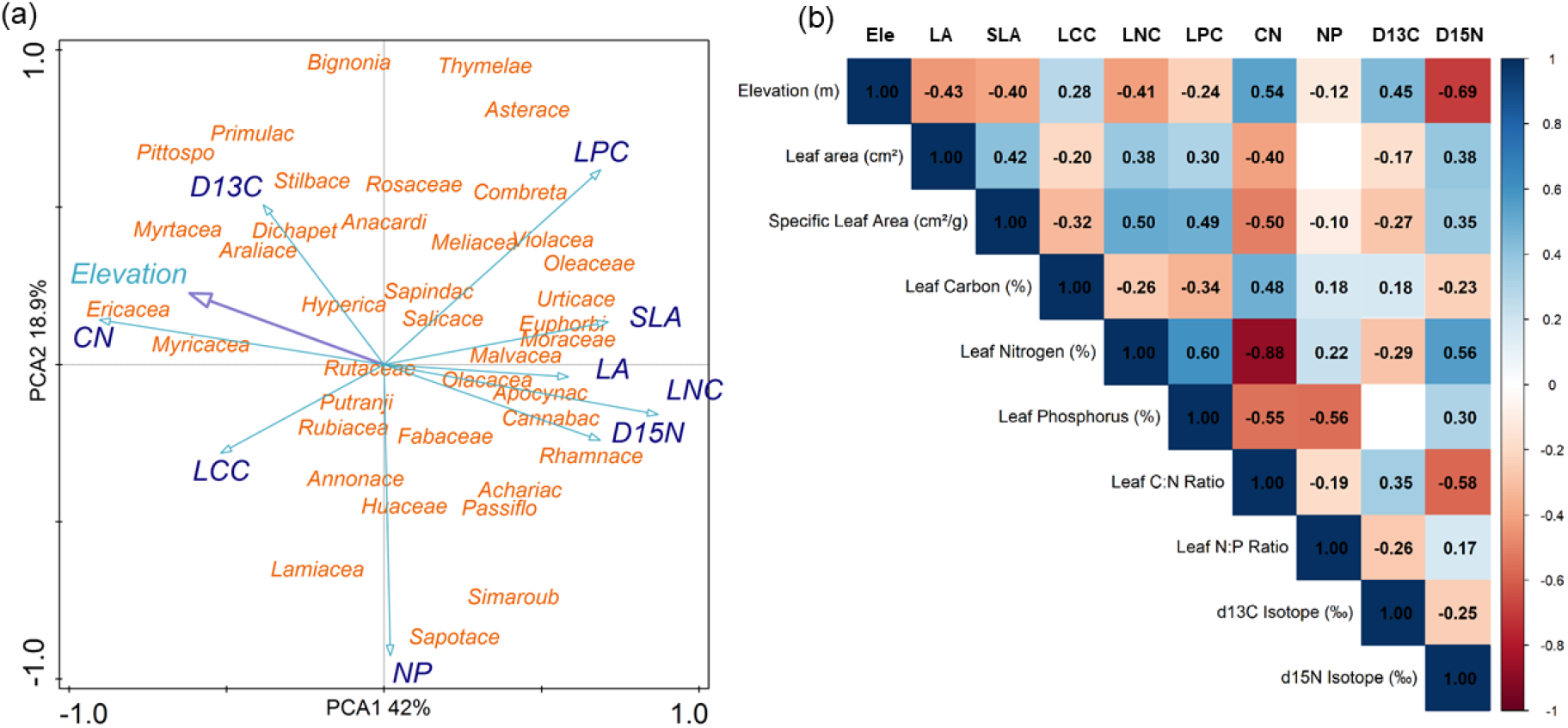
(a) PCA biplot illustrates the relationships between leaf traits and plant families, revealing trait–environment associations and compositional changes with elevation. (b) A correlation matrix showing that elevation is negatively associated with leaf area, nitrogen, phosphorus, and δ^15^N, and positively associated with δ^13^C and the C:N ratio, reflecting shifts in plant functional strategies.

**Figure 5.**
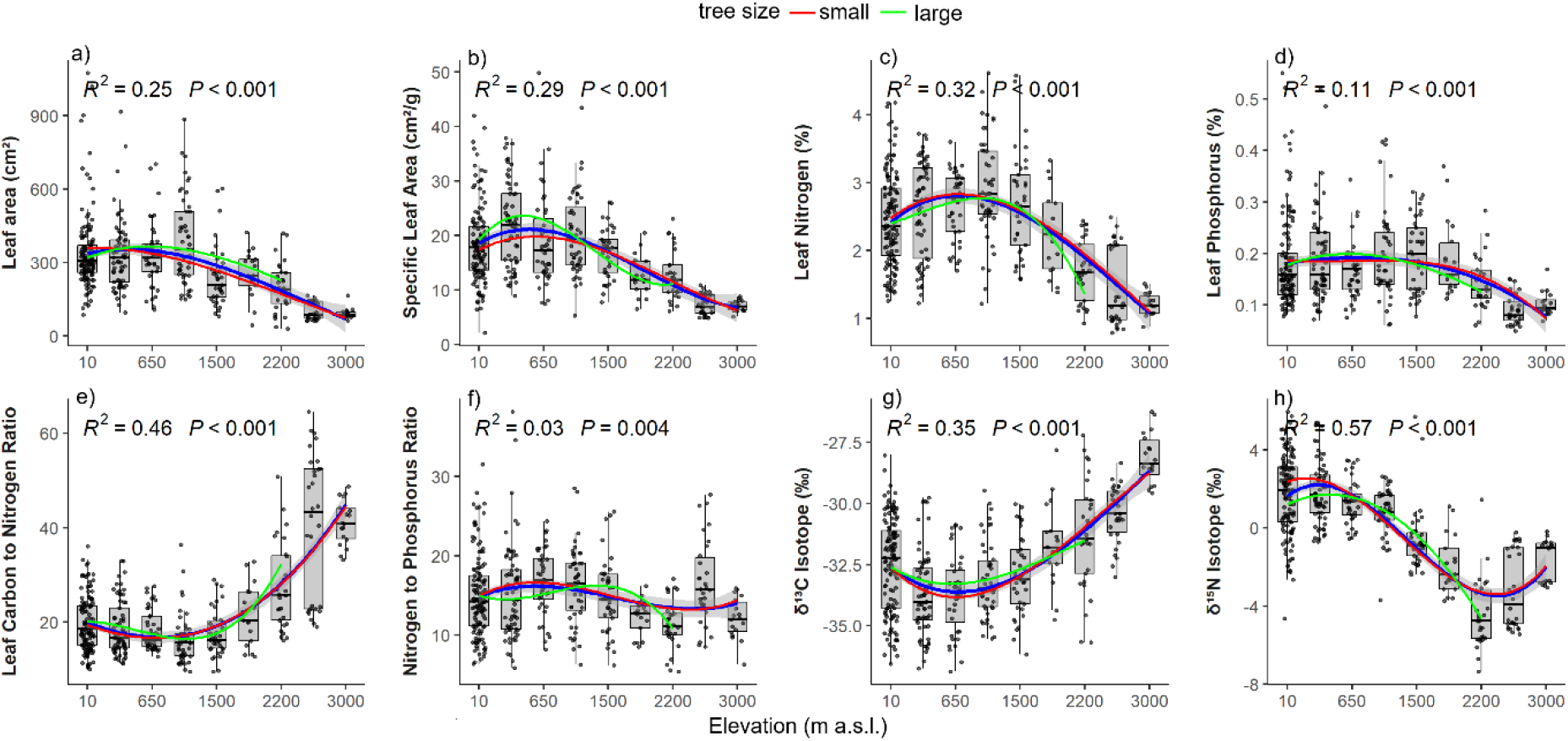
Elevational patterns in leaf functional traits of tropical trees on Mount Cameroon. Relationships between elevation and (a) leaf area, (b) specific leaf area (SLA), (c) leaf nitrogen concentration, (d) leaf phosphorus concentration, (e) leaf carbon-to-nitrogen ratio (C:N), (f) leaf nitrogen-to-phosphorus ratio (N:P), (g) leaf carbon isotope composition (δ^13^C), and (h) leaf nitrogen isotope composition (δ^15^N). Generalized additive model (GAM) fits are shown for all individuals (purple line), small-statured species (red line), and large-statured species (green line). Boxplots represent trait distributions within elevation bands. All relationships are statistically significant (P < 0.001), except for N:P ratio (P = 0.004). Coefficients of determination (R^2^) indicate model fit.

## 3. Results

### 3.1. Environmental and Elevational Gradients

Our study encompassed a 3,200-m elevational gradient along the southwestern flank of Mount Cameroon (Fig. 1a), capturing a broad range of forest types, microclimates, and edaphic conditions (Figs. 2–3). Elevation was strongly and negatively associated with mean annual temperature (R^2^ = 0.95, *P* < 0.001; Fig. 3a), which declined from >25 °C in the lowlands to ∼8 °C at the uppermost elevations with sparse tree cover. Soil moisture displayed a pronounced mid-elevation peak (R^2^ = 0.56, *P* < 0.001; Fig. 3b), mirroring patterns observed for soil organic carbon (R^2^ = 0.29, *P* < 0.001; Fig. 3c), calcium (R^2^ = 0.14, *P* < 0.001; Fig. 3d), and nitrogen (R^2^ = 0.37, *P* < 0.001; Fig. 3e).

Soil phosphorus exhibited a unimodal distribution (R^2^ = 0.21, *P* < 0.001; Fig. 3f), with lowest values from low to mid elevations and a distinct peak around 2,200 m. Stoichiometric ratios varied with elevation: soil C:N declined with altitude up to the montane forest limit (∼2,200 m), followed by an upswing toward the treeline (∼3,200 m; R^2^ = 0.24, *P* < 0.001; Fig. 3g). Soil N:P ratio also followed a unimodal pattern, peaking between 1,500 and 1,600 m (R^2^ = 0.41, *P* < 0.001; Fig. 3h), indicating sharp shifts in nutrient dynamics across ecotones.

### 3.2. Shifts in Leaf Functional Traits

PCA of leaf traits revealed strong trait–environment coordination and phylogenetic clustering along elevation (Fig. 4a). The first two PCA axes captured the dominant patterns across the nine traits, explaining 60.88% of the total variance (Fig. 4a). This axis reflected a gradient from acquisitive strategies, characterized by high leaf area (LA), specific leaf area (SLA), and foliar nitrogen (LNC) and phosphorus (LPC), typically found in lowland rainforest species, to conservative strategies with low SLA, low leaf N and P, and elevated C:N ratios and δ^13^C values, common among high-elevation taxa. Trait means (±standard deviation) were as follows: LA = 295.7 ±180.8 cm^2^, SLA = 17.8 ±9.3 cm^2^/g, LNC = 2.38 ±0.79 %, LPC = 0.17 ±0.08 %, LCC = 45.2 ±2.98 %, C:N = 21.9 ±9.97, N:P = 15.0 ±5.05, δ^13^C = –32.5 ±2.15 ‰, and δ^15^N = 0.32 ±2.63 ‰. High-elevation families, such as Ericaceae, Stilbaceae, and Myricaceae, clustered around high δ^13^C, high C:N, and low SLA and nutrient content. In contrast, lowland families (e.g., Irvinigiaceae, Clusiaceae) exhibited higher SLA, LA, and nutrient concentrations, consistent with fast-growing, resource-acquisitive strategies. Intermediate families, such as Fabaceae and Rosaceae, occupied a broader, ecologically versatile trait space.

### 3.3. Trait Coordination and Stoichiometric Trade-offs Across the Elevational Gradient

Trait correlations mirrored the elevational and environmental gradients, revealing strong coordination among functional, stoichiometric, and isotopic leaf properties (Fig. 4b). Leaf carbon content (LCC) and C:N ratio were positively associated with δ^13^C (r = 0.18 and r = 0.35, P < 0.01), consistent with increased intrinsic water-use efficiency (WUE) under drier or cooler microclimates. This supports the hypothesis that thicker or denser leaves, with higher structural carbon, may exhibit reduced stomatal conductance and thus higher δ^13^C values under stressful conditions. Leaf nitrogen (LNC) and phosphorus content (LPC) were tightly coupled (r = 0.60, P < 0.001), reflecting co-regulation of foliar nutrients involved in photosynthesis and protein synthesis. Both nutrients were strongly and negatively correlated with the C:N ratio (LNC vs. C:N: r = –0.88, P < 0.001; LPC vs. C:N: r = –0.55, P < 0.001), confirming that nutrient-rich leaves exhibit lower stoichiometric C:N ratios and align with acquisitive strategies.

The leaf N:P ratio was negatively associated with LPC (r = –0.56, P < 0.001) and weakly with LNC (r = 0.22, P < 0.05), suggesting that phosphorus availability exerts a stronger control on stoichiometric ratios in this system than nitrogen. δ^15^N exhibited a moderately strong positive correlation with LNC (r = 0.56, P < 0.001) and a weaker correlation with LPC (r = 0.30, P < 0.01), possibly reflecting shifts in nitrogen sources or turnover rates with foliar nutrient availability. However, δ^15^N declined with C:N (r = –0.58, P < 0.001) and elevation (r = –0.69, P < 0.001), supporting the idea that higher-altitude species exhibit more conservative nitrogen cycling or increased mycorrhizal associations that reduce foliar δ^15^N values.

SLA was positively correlated with LNC (r = 0.50, P < 0.001) and LPC (r = 0.49, P < 0.001), and negatively with elevation (r = –0.40, P < 0.001), in line with expectations from the leaf economics spectrum (LES). Similarly, LA showed a positive relationship with SLA (r = 0.42, P < 0.001) and LNC (r = 0.38, P < 0.01), suggesting that large, thin leaves dominate in nutrient-rich lowland forests, while high-elevation communities are characterized by smaller, thicker, nutrient-conservative foliage. Importantly, we observed a negative correlation between LNC and δ^13^C across the gradient (r = –0.29, P < 0.01), consistent with a trade-off between photosynthetic nitrogen efficiency and water-use efficiency in increasingly stressful environments. This pattern reflects the inherent physiological constraint whereby nitrogen-rich, high-photosynthesis leaves often require open stomata and high transpiration rates, resulting in lower WUE and more negative δ^13^C values.

### 3.4. Elevational Trends in Leaf Functional Traits

Leaf traits varied nonlinearly along the 3,000-meter elevational gradient, reflecting shifts in resource-use strategies, physiological constraints, and forest structure (Fig. 5). Specific leaf area (SLA), leaf area (LA), and foliar nitrogen (LNC) and phosphorus (LPC) all showed hump-shaped responses to elevation, peaking at mid-elevations (∼1,000–1,500 m) and declining sharply at higher altitudes. These mid-elevation forests, dominated by tall canopy trees and disturbed by elephants, likely provide optimal conditions for rapid growth and nutrient uptake, combining moderate temperatures, relatively high nutrient availability, and dynamic canopy openings. In contrast, high-elevation forests (>2,000 m) were characterized by smaller, thicker leaves (lower SLA and LA), lower foliar nutrient concentrations, and elevated C:N ratios, consistent with more conservative strategies under colder, windier, and nutrient-constrained conditions. Elevation was negatively correlated with LA (r = –0.43, P < 0.001, Fig. 5a) and SLA (r = –0.40, P < 0.001, Fig. 5b), reflecting anatomical investment in structural leaf tissue at altitude. Stoichiometric ratios showed contrasting responses: the C:N ratio increased steeply with elevation (R^2^ = 0.46, P < 0.001, Fig. 5e), indicating a shift toward greater structural investment and lower nutrient allocation. The N:P ratio displayed a shallow U-shaped pattern, with the lowest values at mid-elevation, where both N and P peaked, again suggesting favorable conditions for balanced nutrient acquisition. Stable isotope signatures further illustrated functional responses to elevation. δ^13^C values increased markedly above 1,000 m (R^2^ = 0.35, P < 0.001, Fig. 5g), suggesting greater intrinsic water-use efficiency (WUE) at higher elevations, where evaporative demand may be high despite lower temperatures. This pattern likely reflects reduced stomatal conductance and thicker leaves that limit CO_2_ diffusion but conserve water. In contrast, δ^15^N declined sharply with elevation (R^2^ = 0.57, P < 0.001, Fig. 5h), indicating either increasing reliance on mycorrhizal nitrogen uptake, lower nitrogen turnover rates, or reduced access to mineral N in cooler soils.

Canopy trees showed higher leaf sizes (LA and SLA) than smaller subcanopy trees, particularly at low to mid elevations. This suggests a more acquisitive trait strategy in canopy dominants, likely reflecting their exposure to higher light and a greater competitive advantage for resource uptake at mid-elevations. However, these differences reverse toward higher elevations, where taller trees showed reduced SLA and LA due to environmental constraints (e.g., temperature, wind, and limited nutrient supply).

### 3.5. Phylogenetically Informed Trait–Environment Relationships

To quantify how environmental factors shape trait variation while accounting for shared evolutionary history, we calculated standardized effect sizes (SES) for 261 trait–environment combinations based on 999 phylogenetically constrained permutations. Of these, 92 associations were statistically significant (|SES| > 2), indicating that observed trait– environment correlations deviated substantially from null expectations under a model of random tip shuffling (Figs. 6 and 7). Traits related to nutrient acquisition and water-use efficiency exhibited the strongest phylogenetic structuring. δ^15^N showed the highest number of significant associations (n = 18), followed by δ^13^C (n = 15), C:N (n = 14), and leaf N (LNC, n = 12). In contrast, traits such as leaf P concentration (LPC) showed little or no phylogenetically structured environmental response. Among the 92 significant associations, 50 were positive (SES > 2) and 42 were negative (SES < –2), suggesting both overdispersion and clustering relative to evolutionary expectation.

**Figure 6.**
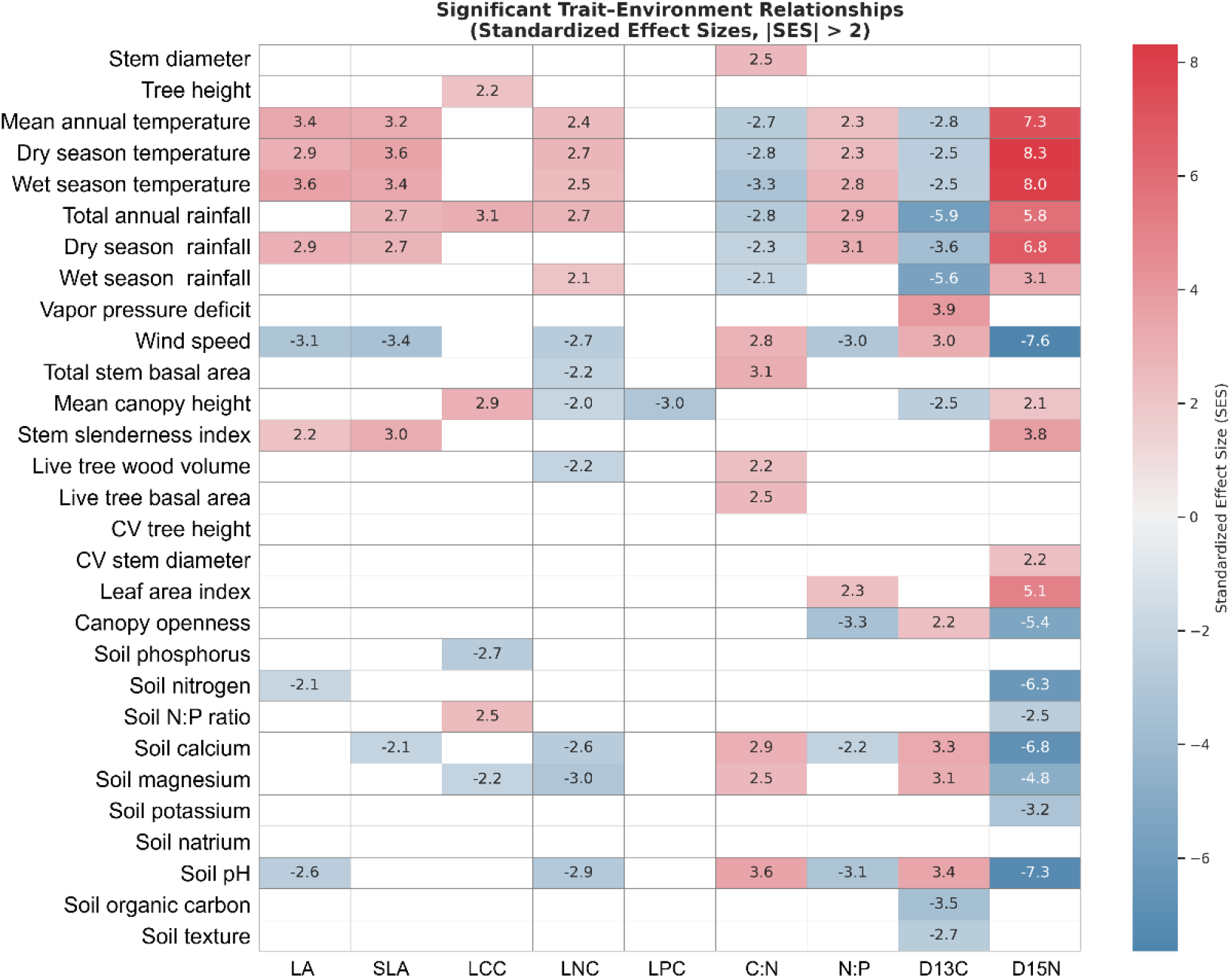
Phylogenetically-structured trait–environment associations across tropical montane tree species. The heatmap shows standardized effect sizes (SES) for the relationships between nine key leaf traits and 29 environmental variables. SES values reflect the deviation from null expectations under phylogenetic constraints. Environmental variables are grouped by tree size variables, climate, forest structure and soil chemistry. Warm colors indicate strong positive associations; cool colors indicate negative associations.

**Figure 7.**
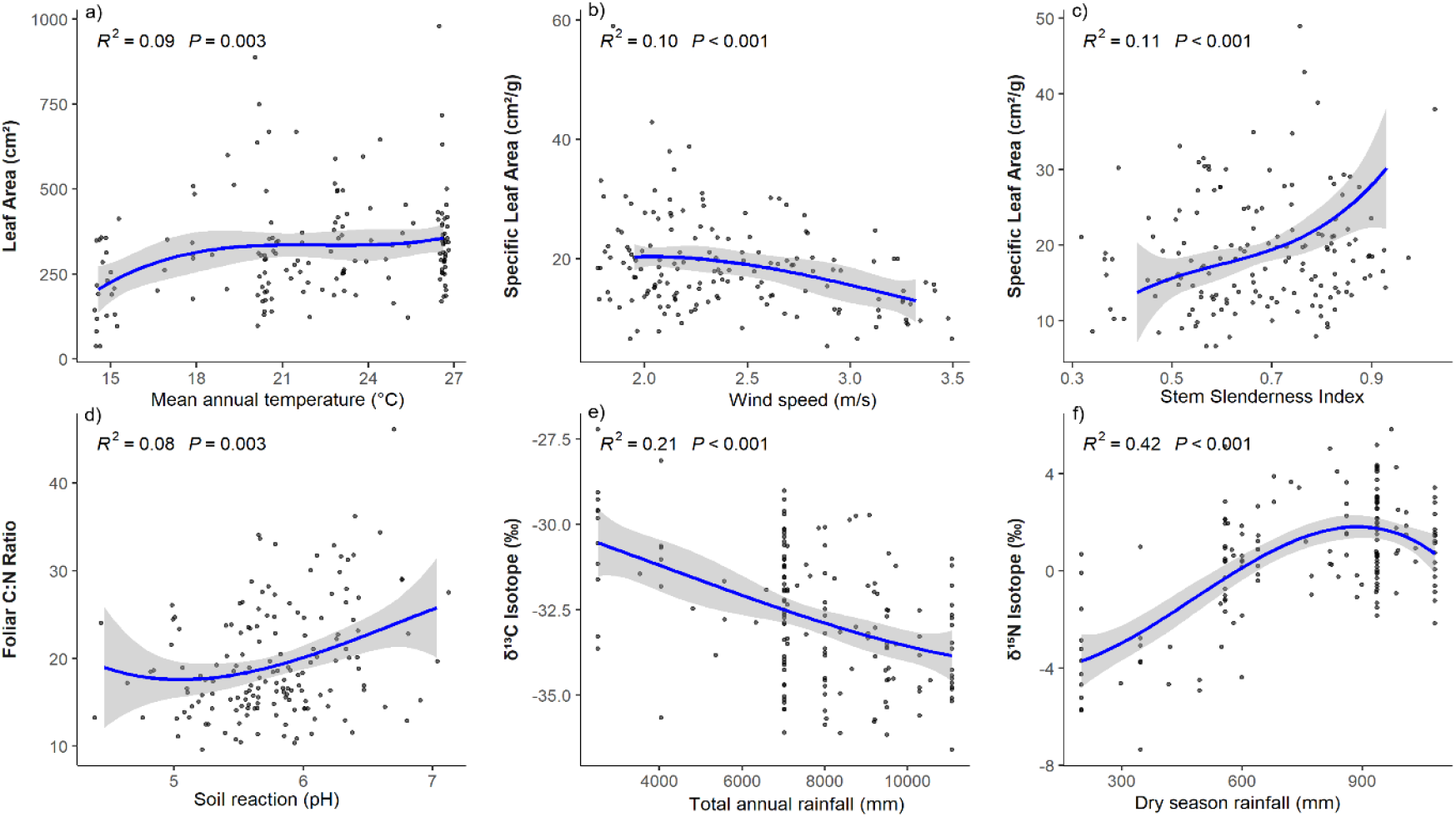
Trait–environment relationships for key foliar traits across the elevational gradient. Panels show associations between (a) leaf area, (b–c) specific leaf area, (d) foliar C:N ratio, (e) δ^13^C, and (f) δ^15^N with selected climatic, edaphic, and structural variables. Lines represent fitted third-order polynomial regressions ±95% confidence intervals.

Climatic variables emerged as the most pervasive environmental filters. δ^13^C declined sharply with rainfall (e.g., SES = –5.9 for total annual rainfall; –5.6 for wet-season rainfall), and increased significantly with wind speed (+3.0), vapor pressure deficit (+3.9), and soil calcium (+3.3), consistent with altered water-use efficiency and stomatal regulation strategies across phylogenetic lineages. δ^15^N increased strongly with mean annual temperature (+7.3), dry-season temperature (+8.3), and wet-season temperature (+8.0), and also responded positively to forest structure metrics, including leaf area index (+5.1) and stem slenderness index (+3.8), while decreasing with wind speed (–7.6) and soil pH (–7.3). These patterns suggest δ^15^N is tightly linked to both thermal regimes and forest physiognomy, potentially reflecting shifts in nitrogen sources and mycorrhizal associations.

The C:N ratio exhibited negative associations with rainfall (e.g., SES = –2.3 with dry-season rainfall) and mean annual temperature (–2.7), and positive associations with several edaphic variables, including soil pH (+3.6), calcium (+2.9), magnesium (+2.5), and soil organic carbon (+3.5), suggesting enhanced structural investment and lower foliar nitrogen in nutrient-poor, acidic environments. N:P ratios also reflected phylogenetically structured environmental filtering, increasing in response to mean annual temperature (+2.3) and dry-season temperature (+2.3), while declining with wet-season rainfall (–5.6) and total rainfall (–2.8).

Soil variables exerted strong but trait-specific filtering. Both δ^13^C and δ^15^N were significantly influenced by soil calcium, magnesium, and pH, but LNC and LPC showed weaker correlations with measured soil nutrients. For example, leaf N declined with soil magnesium (–3.0), while leaf P showed virtually no significant associations, indicating a weak correspondence between soil nutrient availability and foliar N or P concentrations. This suggests that nutrient uptake strategies are highly species-specific or mediated by unmeasured traits such as root architecture or mycorrhizal type.

Forest structure further modulated trait–environment relationships. SLA declined with wind speed (SES = –3.4), possibly reflecting investment in thicker, more durable leaves under mechanical stress. δ^15^N decreased under open canopies (–5.4) and increased with leaf area index (+5.1) and stem basal area (+3.1), indicating that nutrient cycling and nitrogen sources are closely tied to forest canopy complexity and vertical structure. Similarly, δ^13^C increased with canopy openness (+2.2) and decreased with wet-season rainfall (–5.6), further underscoring its role as a sensitive indicator of water-use strategy modulated by both environmental and phylogenetic constraints.

## 4. Discussion

### 4.1. Environmental Drivers of Functional Trait Coordination Along a Tropical Mountain Gradient

Our findings reveal a robust and ecologically coherent shift in leaf functional traits across a 3,200-meter elevational transect on Mount Cameroon, an Afrotropical biodiversity hotspot, highlighting how climate, soils, forest structure, and evolutionary history jointly shape plant strategies in tropical mountain systems. Trait–environment relationships tracked a continuum from acquisitive to conservative leaf syndromes, structured by elevation-dependent changes in temperature, moisture regimes, nutrient availability, and disturbance. Mid-elevation forests (∼1,000–1,500 m), characterized by high productivity, dynamic canopy gaps from elephant disturbance, and moderate climate, harbored species with high SLA, leaf N and P, and low C:N ratios, consistent with fast-growth strategies. In contrast, both lowland rainforests (500– 1000 m) and upper montane habitats (>2,000 m) supported more stress-tolerant species with smaller, tougher leaves, lower nutrient content, and higher δ^13^C and C:N values, shaped respectively by wet-season waterlogging or cold and fire-driven nutrient stress. These nonlinear patterns reinforce recent calls to move beyond linear or monotonic expectations in trait– environment relationships across tropical elevation gradients (Fyllas et al., 2017; Oliveras et al., 2020).

Stable isotopes (δ^13^C, δ^15^N) and stoichiometric traits (C:N, N:P) emerged as particularly sensitive indicators of environmental filtering. Their variation captured integrated responses to microclimate, moisture, soil nutrient supply, and mycorrhizal strategies, and their strong phylogenetic signal underscores the role of evolutionary history in shaping trait–environment associations. The combined influence of climate, fire, hydrology, and canopy architecture on trait distributions, modulated by taxonomic lineage, supports a model of multidimensional environmental filtering where biotic and abiotic constraints act in concert. Notably, our results diverge from global expectations linking leaf nutrient content tightly to soil fertility (e.g., Ordoñez et al., 2009), instead emphasizing the importance of hydrological context, leaching, and disturbance in decoupling foliar stoichiometry from total soil nutrient pools in hyper-humid tropical systems.

### 4.2. Trait Shifts Along Elevation: A Move Toward Conservative Strategies

The observed decline in specific leaf area (SLA), leaf area (LA), and nutrient concentrations (N and P), along with increasing carbon-to-nitrogen (C:N) ratios and δ^13^C values with elevation, indicates a shift toward more conservative resource-use strategies under cooler, nutrient-poor, and potentially water-limited conditions. These patterns are consistent with findings from Andean and Asian montane systems (Homeier et al., 2021; Helsen et al., 2023), where declining temperatures, nutrient availability, and increasing evaporative demand jointly select for thicker, denser leaves with longer lifespans. Oliveras et al. (2020) similarly reported consistent elevation-related trait shifts across three tropical environmental gradients in South America and Africa, where leaf structure and chemistry co-varied predictably with climate and soil properties. Canopy trees, particularly in lowland forests, consistently exhibited higher SLA than subcanopy individuals, likely reflecting access to deeper nutrient pools and a stronger competitive advantage for light. However, these differences diminished at higher elevations, where temperature and nutrient constraints override vertical stratification, echoing patterns reported from other tropical mountains (Kitayama & Aiba, 2002; Moser et al., 2007).

Foliar δ^13^C values increased markedly above 1000 m, suggesting enhanced intrinsic water-use efficiency (WUE), likely due to reduced stomatal conductance (Cernusak et al., 2007; Cornwell et al., 2018). The parallel increase in foliar carbon content and C:N ratio further supports a shift toward structural investment over nutrient acquisition. As leaves become thicker and denser with elevation, reflected in declining specific leaf area (SLA), they tend to exhibit reduced nutrient concentrations and higher δ^13^C values. This pattern likely reflects a shift in resource allocation strategies under cold, dry, and irradiated conditions. Thicker leaves, while conferring greater durability and resistance to desiccation or photodamage, often reduce mesophyll conductance, thereby limiting CO_2_ diffusion to the chloroplasts and lowering photosynthetic capacity per unit nitrogen (Wright et al., 2001; Cornwell et al., 2018). The associated increase in δ^13^C reflects lower intercellular CO_2_ concentrations, thereby enhancing WUE. Accordingly, we observed a negative correlation between leaf nitrogen content and δ^13^C across the gradient, consistent with a trade-off between photosynthetic nitrogen efficiency and water-use efficiency in increasingly stressful environments. The inverse relationship between LNC and δ^13^C thus highlights the coordinated adjustment of structural, nutritional, and physiological traits in response to the shifting balance between carbon acquisition and stress tolerance along the elevation gradient. Similar trait shifts have been documented in montane forests of Taiwan (Helsen et al., 2023) and in global meta-analyses across elevational gradients (Fyllas et al., 2017)

### 4.3. Stable Carbon Isotopes Trace Plant Responses to Moisture and Elevational Stress

Stable isotope data provided further insight into plant ecophysiology. Foliar δ^13^C values increased with elevation, indicating greater WUE or diffusion limitation of CO_2_ under cold, low-pressure conditions, consistent with denser leaf structures and lower SLA (Wright et al., 2001; Cornwell et al., 2018). The steep decline in δ^13^C values observed along the elevational gradient from approximately 1000 to 3000 m deviates markedly from the global average of ∼1.1‰ per kilometer typically reported for humid, well-ventilated sites (Körner et al., 1991). Instead, the slope in our dataset is substantially steeper, likely reflecting the combined influence of changing moisture regimes, forest structure, and atmospheric conditions. Notably, total annual precipitation drops sharply from 10,000–12,000 mm at low elevations (300–1100 m) to less than 5000 mm between 1100 and 1600 m, and further declines to below 500 mm in the afromontane zone above 2400 m, where *Agarista salicifolia* and *Myrica arborea* trees occur in scattered savanna formations (Dolezal et al., 2022). Above 1600 m, montane forests experience a pronounced dry season from December to March, which likely contributes to reduced stomatal conductance and elevated δ^13^C values at higher elevations. In contrast, the very negative δ^13^C values recorded at lower elevations, approaching –32‰, may result from consistently high humidity and low light availability under a near-permanent cloud layer in the lowland rainforest, as well as the influence of soil-respired CO_2_ enriched in ^12^C. Such conditions elevate internal CO_2_ concentrations, enhancing discrimination against ^13^C. Collectively, these findings indicate a non-linear, moisture-sensitive response of foliar δ^13^C to elevation, characterized by a transition from saturated, shaded lowland forests to seasonally dry montane zones and arid Afroalpine environments.

### 4.4. Waterlogging Effects on Foliar δ^13^C in Lowland Rainforests

The very negative foliar δ^13^C values observed below 1000 m may, in part, reflect the physiological effects of waterlogging associated with extremely high annual rainfall, exceeding 10,000–12,000 mm, in these lowland rainforests. Prolonged saturation of soils can lead to anoxic conditions, impairing root function and reducing both water and nutrient uptake. This in turn can constrain photosynthetic capacity more strongly than stomatal conductance, resulting in elevated intercellular CO_2_ concentrations (Ci) and increased discrimination against ^13^C. The consequence is a shift toward more negative δ^13^C values in leaf tissues. Additionally, waterlogged soils often emit respired CO_2_ derived from anaerobic decomposition, which carries a depleted ^13^C signature. In humid, poorly ventilated forest understoreys, re-fixation of this CO_2_ may further reduce foliar δ^13^C. These processes, combined with low vapor pressure deficit (VPD) and deep canopy shading, mirror the physiological effects of cloud immersion and reinforce the observed δ^13^C depletion at low elevations. Thus, the extremely moist conditions of the lowland zone likely contribute significantly to the lower δ^13^C values, independent of elevation per se, and should be considered when interpreting carbon isotope patterns across tropical mountain gradients.

### 4.5. Nitrogen Isotope Signatures Trace Environmental Filtering and Symbiotic Strategies

Foliar δ^15^N values declined markedly with elevation across the Mount Cameroon gradient, ranging from approximately +5‰ in lowland forests to −5‰ at higher elevations. This trend reflects a shift from mineral nitrogen acquisition toward organic and mycorrhiza-mediated uptake in cooler, less biologically active environments, in line with global patterns (Hobbie and Högberg, 2012; Craine et al., 2015). According to known isotopic signatures, δ^15^N values around 0‰ typically indicate nitrogen fixation, values near −2‰ reflect arbuscular mycorrhizal (AM) associations, and more negative values (–4‰ to –6‰) are characteristic of ectomycorrhizal (EM) and ericoid mycorrhizal (ERM) strategies (Hobbie and Högberg, 2012). Our dataset supports these distinctions: species from Fabaceae showed relatively enriched δ^15^N values in lowland sites, consistent with nitrogen-fixation and AM-mediated uptake under high rainfall and leaching. In contrast, at the upper elevational limit, *Agarista salicifolia* (Ericaceae) exhibited a notably depleted δ^15^N value of –3.2‰, suggesting ERM-mediated nitrogen acquisition in highly organic, N-poor soils typical of high montane ecosystems. This agrees with the observation by Hobbie and Högberg (2012) that ERM plants acquire nitrogen from recalcitrant organic sources, leading to strong isotopic discrimination and depleted foliar δ^15^N.

Additionally, a bimodal distribution in δ^15^N values was observed. The first enrichment peak (0–1000 m) coincides with intense precipitation and nitrogen leaching on the southwestern slope, which favors nitrogen-fixing taxa, such as Fabaceae. The second peak occurred in upper montane forests (2,200–2,500 m), where frequent dry-season fires likely induce nitrogen losses through volatilization (Doležal et al., 2022). Here, the dominance of *Myrica arborea* (Myricaceae), another nitrogen-fixing species, underscores the role of disturbance-driven nutrient scarcity in shaping nitrogen acquisition strategies. In both cases, enriched δ^15^N signals correspond to disturbance-related nitrogen scarcity and nitrogen fixation capability. Together, these findings suggest that δ^15^N variation in tropical trees is jointly driven by environmental filtering (rainfall, temperature, fire) and phylogenetically constrained symbiotic strategies (e.g., N-fixation, AM, EM, ERM), echoing patterns reported by Hobbie and Högberg (2012) and reinforcing the functional link between nitrogen isotopes, mycorrhizal type, and plant adaptation across elevation.

Foliar δ^15^N showed a strong positive correlation with dry-season rainfall (R^2^ = 0.42, *P* < 0.001; Fig. 7), suggesting that wetter dry seasons promote nitrogen losses via leaching or denitrification, enriching the δ^15^N of residual soil nitrogen. This isotopic signal corresponds spatially with the lowland rainforest zone on Mt. Cameroon, where tree growth is paradoxically constrained during the peak wet season due to anoxia and low light conditions under high cloud cover (Plavcová et al., 2024). In these forests, tree radial growth often peaks during the dry season, when soils are aerated but remain sufficiently moist to permit active xylem expansion. In contrast, upper montane trees experience growth limitation during the dry season due to water scarcity. The elevated δ^15^N values in wetter lowland forests may thus reflect both increased nitrogen loss and the prevalence of nitrogen-fixing species that dominate under such leached, anaerobic conditions. This co-occurrence of enriched δ^15^N and growth suppression during the wettest periods highlights the complex coupling between nitrogen cycling, water availability, and growth phenology in Afrotropical forests.

### 4.6. Divergence from Global Patterns: Decoupling of Leaf Nutrients from Soil Fertility

Our results contrast markedly with global-scale findings by Ordoñez et al. (2009), who reported that SLA, LNC, and LPC generally increased with soil fertility across ecosystems worldwide. In our Afrotropical elevational gradient on Mt. Cameroon, we found no consistent relationship between foliar N or P concentrations and corresponding soil nutrient pools. This decoupling likely reflects the overriding influence of extreme rainfall and leaching in low-elevation tropical rainforests (650 m), where soils are heavily weathered, organic matter is low, and nutrient retention is poor. Such conditions suppress mineralization and reduce nutrient bioavailability despite high productivity and biomass.

Indeed, the lowest soil and foliar N and P values were observed in these hyper-humid lowland forests, which receive up to 10,000 mm of rainfall annually. This stands in contrast to global patterns where soil nutrient levels explain substantial variation in foliar traits (cf. Ordoñez et al., 2009; Wright et al., 2004). Only in mid-elevation forests between 1500 and 1800 m, subject to frequent disturbance by forest elephants, did we observe concurrent peaks in both soil and leaf nutrient concentrations. These disturbed zones appear to maintain more dynamic nutrient turnover through bioturbation, canopy gap formation, and enhanced microbial activity, reinforcing localized nutrient availability and uptake.

This pattern highlights how edaphic control of leaf functional traits may be contingent on hydrological context and disturbance regimes. In hyper-humid systems, leaching and redox constraints decouple foliar stoichiometry from soil total nutrient pools (e.g., Vitousek et al., 2010; Townsend et al., 2007), while in montane systems, moderate disturbance and improved drainage can enhance nutrient accessibility despite lower temperatures. These findings underscore the limitations of applying generalized global trait–soil models to hyperdiverse, moisture-saturated ecosystems and support calls to incorporate regional climatic and ecological dynamics into predictive trait-based models (e.g., Fyllas et al., 2017; Oliveras et al., 2020).

### 4.7. Phosphorus Availability is Shaped by Hydrology and Fire, Not Elevation Alone

Contrary to expectations based on the leaf economic spectrum (LES), our results reveal that phosphorus limitation is most pronounced at lower elevations, rather than at high-elevation sites, where colder temperatures and slower nutrient cycling are typically thought to constrain P availability (Vitousek et al., 2010 and references therein). Both foliar and soil phosphorus concentrations were lowest between 500 and 1000 m elevation, within the hyper-humid rainforest belt receiving up to 10,000 mm of annual rainfall (řeháková et al., 2022). These lowland soils are often organic-poor, poorly drained, and subject to temporal waterlogging, creating anoxic conditions that suppress mineralization and promote phosphorus fixation through redox-sensitive sorption processes. In contrast, mid- to high-elevation soils (2200– 2400 m) exhibited notably higher available phosphorus concentrations, coinciding with montane forest and Afroalpine savanna zones that experience regular dry-season fires during January (Doležal et al., 2022). These fire events can enhance soil phosphorus availability through combustion of organic material, ash deposition, and altered microbial turnover, contributing to transient but ecologically significant peaks in soil P at higher elevations (Hogue & Inglett, 2012; Veldhuis et al., 2016).

Interestingly, while soil P peaks at higher elevations, foliar P content displays the opposite pattern, declining consistently with elevation in line with LES predictions. This divergence suggests that low leaf P at high elevation is not driven by reduced soil P availability, but rather by physiological or morphological constraints on nutrient uptake, transport, or demand. Cold temperatures may reduce root activity and phosphorus mobility, while shorter growing seasons and conservative life-history strategies may reduce foliar nutrient investment despite adequate supply in the soil. Conversely, in the lowlands, the combination of high foliar P demand and low P bioavailability in saturated, leached, and redox-active soils likely intensifies actual phosphorus limitation. Together, these findings reveal that soil and plant phosphorus dynamics are decoupled along elevation gradients and shaped by the interplay of hydrological context, disturbance regimes, and plant functional strategies.

### 4.8. Stoichiometric Shifts and Nutrient Limitation Along Elevation

Elevational patterns in both soil and foliar nutrient stoichiometry revealed significant nonlinear shifts in ecosystem nutrient dynamics along the Mount Cameroon gradient. The soil nitrogen-to-phosphorus (N:P) ratio exhibited a pronounced hump-shaped response to elevation (R^2^ = 0.41, *P* < 0.001), peaking in mid-elevation forests (∼1500 m) that are regularly disturbed by forest elephants. These open-canopy sites likely support enhanced microbial activity, nutrient cycling, and mineralization, driven by increased light availability, accelerated litter decomposition, and nutrient redistribution through dung deposition and trampling. In contrast, foliar N:P ratios showed a much weaker elevational trend (R^2^ = 0.03, *P* = 0.004), though they too peaked modestly in the same mid-elevation zone. This partial decoupling between soil and foliar stoichiometry highlights the influence of additional factors, including species identity, functional strategy, and mycorrhizal associations, on leaf-level nutrient ratios.

Our observed foliar N:P ratios largely fell within the intermediate range (10–20), suggesting co-limitation by N and P. However, higher values at mid-elevation may indicate increasing P limitation in sites with relatively high N availability. These patterns are broadly consistent with the global synthesis by Güsewell (2004), who demonstrated that foliar N:P ratios reflect both nutrient supply and a species’ ecological strategies, with higher N:P values being more common in stress-tolerant or graminoid species. They also align with findings from Townsend et al. (2007), who emphasized that while tropical forests are often considered N-rich and P-limited, foliar N:P ratios are not tightly constrained across tropical ecosystems, largely due to species-level variability and local environmental heterogeneity. Similarly, Soethe et al. (2008) found elevated foliar N:P at mid-elevations in Ecuatorian montane forests, where litter decomposition was more active and nutrient availability more favorable than at higher altitudes.

### 4.9. Phylogenetically Structured Environmental Filtering

Our results indicate that environmental filtering of functional traits in tropical forest communities is profoundly shaped by phylogenetic history (Cavender-Bares et al., 2009; Swenson and Enquist, 2009). Traits related to nutrient acquisition and physiological regulation, particularly δ^15^N, δ^13^C, and C:N, demonstrated the strongest departures from phylogenetically randomized null expectations, indicating that both climate and soil gradients select for conserved trait syndromes within evolutionary lineages (Cornwell & Ackerly, 2009; Laughlin & Messier, 2015). Climatic factors were key drivers of trait differentiation. The strong negative SES values for δ^13^C in wetter environments and positive values with wind stress suggest that lineages differ in intrinsic stomatal behavior and water-use efficiency, shaped by both habitat moisture and mechanical exposure (Farquhar et al., 1989; Cernusak et al., 2007; Cornwell et al., 2018). Similarly, the consistent positive associations of δ^15^N with temperature and LAI may reflect evolutionary shifts in nitrogen source pools, uptake mechanisms, or mycorrhizal symbioses in warmer and less disturbed environments (Högberg, 1997; Craine et al., 2015; Mayor et al., 2014).

Soil properties exerted clear but trait-specific constraints. The positive SES values of C:N with base saturation and pH imply that nutrient allocation strategies are tightly linked to edaphic fertility and acidity, with conservative nutrient use prevailing in lineages associated with richer soils (Townsend et al., 2007; Porder & Hilley, 2011). Increasing base saturation and reduced nitrogen availability at higher pH may constrain nitrogen uptake relative to carbon assimilation. As soil pH rises, ammonium is increasingly converted to nitrate, which is more mobile and susceptible to leaching in humid tropical systems (Vitousek & Sanford, 1986). Additionally, enhanced microbial activity under near-neutral pH can promote nitrogen immobilization in microbial biomass, further reducing nitrogen availability to plants (Güsewell, 2004). Base-rich soils may also induce nutrient imbalances or competition between cations (e.g. Ca^2^_+_, Mg^2^_+_, K_+_) and ammonium, limiting nitrogen uptake (Townsend et al., 2007). The resulting increase in foliar C:N reflects a shift toward nutrient-conservative strategies, emphasizing structural over metabolic investment under nutrient-limiting conditions. Similar patterns have been reported from base-rich montane forests and fire-affected soils (Oliveras et al., 2020; Santiago et al., 2017), supporting the interpretation that foliar stoichiometry integrates both soil chemistry and functional adaptation to resource constraints.

By contrast, traits such as P showed weak or absent phylogenetic signal, suggesting higher plasticity or lower evolutionary constraint (Poorter et al., 2009; Baraloto et al., 2010). Forest structure also shaped trait patterns in phylogenetically non-random ways. Increases in C:N and δ^15^N with openness, and declines in SLA with wind exposure, point to the role of vertical canopy architecture and microclimatic variation in shaping resource-use and structural traits (Kitajima & Poorter, 2010; Markesteijn et al., 2011). These relationships reflect how mechanical stress and light availability interact with evolutionary legacy to constrain functional adaptation (Asner et al., 2018; Givnish, 2002).

Functional differentiation and community assembly patterns revealed further insight into evolutionary constraints. PCA showed distinct trait syndromes and strong family-level clustering. High-elevation lineages, such as Ericaceae, Myrtaceae, and Myricaceae, were associated with conservative strategies including high δ^13^C and C:N, and low SLA and foliar nutrient content, reflecting adaptations to cold, nutrient-poor, or fire-prone environments. In contrast, lowland families such as Irvingiaceae and Clusiaceae exhibited acquisitive traits associated with rapid growth under warm, moist, and fertile conditions, aligning with global LES expectations (Wright et al., 2004; Díaz et al., 2016). Some families, such as Fabaceae and Rosaceae, occupied broader functional spaces. Fabaceae in particular maintained elevated δ^15^N values across the gradient, consistent with their N-fixing capacity and resilience under both leaching and fire-prone regimes. This taxonomic control over trait expression reinforces findings by Oliveras et al. (2020), who demonstrated that family identity often explains more variance in plant traits than site-level environmental variation.

Taken together, these findings support a model of multidimensional environmental filtering where functional traits are shaped not only by climate and soils but also by lineage-specific evolutionary history and disturbance regimes. Acquisitive trait syndromes (e.g., high SLA, N, P) are favored in warm, nutrient-rich, open forests, while conservative syndromes (e.g., high C:N, δ^13^C) dominate in cooler, less fertile, or more structurally complex habitats. The strength and direction of SES patterns, combined with phylogenetic clustering in multivariate trait space, highlight the importance of integrating taxonomic and evolutionary information when interpreting trait–environment relationships and forecasting tropical forest responses to global change (Fyllas et al., 2017; Swenson & Enquist, 2009).

## 5. Synthesis and Implications for Tropical Trait-Based Ecology

In sum, the Mount Cameroon elevational gradient reveals strong and coordinated shifts in plant functional traits that reflect both adaptive responses to local environmental stressors and deep phylogenetic legacies. Trait–environment relationships, especially those involving SLA, C:N, δ^13^C, and δ^15^N, aligned with resource-use trade-offs that were modulated by temperature, precipitation, fire regimes, and soil chemistry, but also structured by lineage-specific constraints and strategies. Mid-elevation forests emerged as functional hotspots, combining high productivity, canopy disturbance, and nutrient availability to support acquisitive leaf traits. In contrast, lowland and highland forests converged on conservative strategies through different mechanisms, waterlogging and nutrient leaching in the former, cold stress and fire-driven nutrient losses in the latter.

These results reinforce the global utility of trait-based approaches in tropical forest ecology, especially when combined with stable isotope and stoichiometric markers that integrate physiological function and environmental history. As climate change accelerates, altering temperature regimes, rainfall seasonality, and fire frequency across tropical mountains (Fyllas et al., 2017; Helsen et al., 2023), the trait strategies documented here can serve as predictive tools for understanding shifts in community composition, productivity, and biogeochemical cycling. Our findings underscore the need for regionally grounded, mechanistically informed trait-based models that account for hydrology, disturbance, and evolutionary constraints when forecasting forest responses to global environmental change.

## Author contributions

JD conceived the ideas and designed the methodology; JD, VB and KK collected the data; JD and KK analysed the data; JD, AR and JG led the writing of the manuscript. All authors contributed critically to the drafts and gave final approval for publication.

## Data Availability Statement

Data available from the Zenodo repository doi: XXX (Dolezal, 2025).

